# Birthweight, Type 2 Diabetes and Cardiovascular Disease: Addressing the Barker Hypothesis with Mendelian randomization

**DOI:** 10.1101/208629

**Authors:** Daniela Zanetti, Emmi Tikkanen, Stefan Gustafsson, James Rush Priest, Stephen Burgess, Erik Ingelsson

**Author notes:** **Address for Correspondence:** Erik Ingelsson, MD, PhD, FAHA, 300 Pasteur Dr, mail code: 5773; Stanford, CA 94305; USA.

## Abstract

**Background:** Low birthweight (BW) has been associated with a higher risk of hypertension, type 2 diabetes (T2D) and cardiovascular disease (CVD) in epidemiological studies. The Barker hypothesis posits that intrauterine growth restriction resulting in lower BW is causal for these diseases, but causality and mechanisms are difficult to infer from observational studies. Mendelian randomization (MR) is a new tool to address this important question.

**Methods:** We performed regression analyses to assess associations of self-reported BW with CVD and T2D in 237,631 individuals from the UK Biobank, a large population-based cohort study aged 40-69 years recruited across UK in 2006-2010. Further, we assessed the causal relationship of such associations using the two-sample MR approach, estimating the causal effect by contrasting the SNP effects on the exposure with the SNP effects on the outcome using independent publicly available genome-wide association datasets.

**Results:** In the observational analyses, BW showed strong inverse associations with systolic and diastolic blood pressure (*β*, −0.83 and −0.26; per raw unit in outcomes and SD change in BW; 95% CI, −0.90, −0.75 and −0.31, −0.22, respectively), T2D (odds ratio [OR], 0.83; 95% CI, 0.79, 0.87), lipid-lowering treatment (OR, 0.84; 95% CI, 0.81, 0.86) and CAD (hazard ratio [HR] 0.85; 95% CI, 0.78, 0.94); while the associations with adult body mass index (BMI) and body fat (*β*, 0.04 and 0.02; per SD change in outcomes and BW; 95% CI, 0.03, 0.04 and 0.01, 0.02, respectively) were positive. The MR analyses indicated inverse causal associations of BW with low density lipoprotein cholesterol, 2-hour glucose, CAD and T2D, and positive causal association with BMI; but no associations with blood pressure. Sensitivity analyses and robust MR methods provided consistent results and indicated no horizontal pleiotropy.

**Conclusion:** Our study indicates that lower BW is causally and directly related with increased susceptibility to CAD and T2D in adulthood. This causal relationship is not mediated by adult obesity or hypertension.

## Introduction

The association between low birthweight (BW) and increased risk of coronary artery disease (CAD) in adult life was first demonstrated by the British epidemiologist David Barker in a landmark paper in *the Lancet* in 1989^1^. This observation was later extended using a longitudinal cohort study of 8,760 participants with growth trajectories during childhood^2^. In this study, individuals with a low BW increased their weight rapidly after two years of age, and had increased risk of insulin resistance and CAD in adult life. In 1992, Barker proposed that these relationships could be explained by what he called the “*Thrifty phenotype hypothesis*”^3^ attributing the association between poor fetal and infant growth and subsequent increased cardiovascular risk to arise from a compensatory response to nutritional deprivation in early life, resulting in permanent changes in glucose-insulin metabolism and somatic growth lasting into adulthood. Decreased insulin secretion and increased insulin resistance in combination with effects of obesity, ageing and physical inactivity are the most important factors leading to type 2 diabetes (T2D)^3^, but they are also independent risk factors for CAD, stroke, and hypertension^4^.

Still, it is not yet clear whether BW plays a causal role in the development of these outcomes as posited in the “Barker Hypothesis”; or if other phenomena, such as confounding factors (maternal smoking, socioeconomics level, ethnicity) have resulted in spurious associations in previous observational studies. We wanted to investigate causal mechanisms using the Mendelian randomization (MR) approach. This method has the ability to infer a causal relationship between a risk factor and a disease, using genetic markers as a proxy for a modifiable exposure. Two smaller prior MR studies indicated a causal association between low BW and T2D^5^, but not with lipids or CAD^6^. However, these studies were hampered by weak instrumental variables including only five and seven single nucleotide polymorphisms (SNPs), respectively; resulting in limited statistical power. Furthermore, these studies did not address the relationship of BW with other important cardiovascular diseases and risk factors, including atrial fibrillation (AF), ischemic stroke (IS), blood pressure, body mass index (BMI), waist-to-hip ratio (WHR), high density lipoproteins (HDL), low density lipoprotein (LDL), triglycerides (TG), 2-hour glucose, fasting glucose, and fasting insulin.

The aims of the present study were to: 1) describe the relationships of self-reported BW to several cardiovascular traits in 237,631 participants of the UK Biobank (UKB); and 2) delineate any causal relationships between BW and CAD, AF, IS and T2D, and risk factors for these diseases (systolic and diastolic blood pressure [SBP and DBP], BMI, WHR, HDL, LDL, TG, 2-hour glucose, fasting glucose, and fasting insulin) by two-sample MR analysis using summary statistics from the largest available genome-wide association study (GWAS) meta-analyses.

## Methods

### Study sample

The UKB is a longitudinal cohort study of over 500,000 individuals aged 40-69 years initiated in the United Kingdom (UK) in 2006-2010^7^. We included 237,631 participants that knew their BW, to focus on the linear effects of birth weight we limited analysis to individuals reporting birth weights to be within 2.5 kg and 4.5 kg, and excluded individuals with CV prior enrollment (Supplemental Table I Online Data Supplement for details). We used UKB for our observational analyses, as well as to perform a GWAS of SBP and DBP (as publically available summary statistics were adjusted for BMI) to create an instrumental variable (IV) for the MR analyses. Cardiovascular outcomes for observational studies were defined using the International Classification of Diseases (ICD) codes (details in the Online Data Supplement). The exposure of interest was self-reported BW.

We used publicly available GWAS summary statistic of BW^8^ as exposure; and of CAD^9^, AF^10^, IS^11^, SBP and DBP adjusted for BMI^12^, BMI^13^, WHR^14^, HDL, LDL, TG^15^, T2D^16^, 2-hour glucose^17^, fasting glucose, and fasting insulin^18^ as outcomes. Details on the GWAS consortia, number of samples, proportion of variance explained and statistical power for MR analysis are presented in Table 1.

**Table 1.** Description of data used and statistical power for Mendelian randomization analyses.

| Phenotype | Consortium | N samples | Variants in the IV2 | Variance explained (%) | Effect in UKB | Power for observed association (%) | Power for fixed standardized effect | Units | Publication |
| --- | --- | --- | --- | --- | --- | --- | --- | --- | --- |
| BW | EGG | 143,677 |  |  |  |  |  | SD (kg/m<sup>2</sup>) | Horikoshi et al., 2016 |
| CAD | CARDioGRAMplusC4D | 184,305 | 45 | 0.022 | 0.854 | 99 | 100 | log odds | Nikpay et al., 2015 |
| AF | AFGen | 133,073 | 39 | 0.020 | 1.179 | 84 | 90 | log odds | Christophersen et al., 2017 |
| IS | ISGC | 435,001 | 45 | 0.022 | 0.881 | 94 | 99 | log odds | Pulit et al., 2016 |
| SBP | UKB | 337,229 | 33 | 0.022 | -0.042 | 95 | 100 | mmHg | Sudlow et al., 2015 |
| DBP | UKB | 337,235 | 33 | 0.022 | -0.025 | 58 | 100 | mmHg |  |
| SBP | ICBP | 201,529 | 34 | 0.020 | -0.042 | 76 | 100 | mmHg | Ehret et al., 2016 |
| DBP | ICBP | 201,529 | 34 | 0.020 | -0.025 | 35 | 100 | mmHg |  |
| BMI | GIANT | 339,224 | 38 | 0.020 | 0.041 | 92 | 100 | SD (kg/m<sup>2</sup>) | Locke et al., 2015 |
| WHR | GIANT | 210,082 | 38 | 0.020 | 0.003 | 4 | 100 | SD | Shungin et al., 2015 |
| HDL | GLGC | 187,167 | 38 | 0.020 | NA | NA | 100 | SD (mg/dL) | Willer CJ et al., 2013 |
| LDL | GLGC | 173,082 | 38 | 0.020 | NA | NA | 100 | SD (mg/dL) |  |
| TG | GLGC | 177,861 | 37 | 0.020 | NA | NA | 100 | SD (mg/dL) |  |
| T2D | DIAGRAM | 149,821 | 17 | 0.012 | 0.832 | 92 | 91 | log odds | Morris et al., 2012 |
| 2hr glucose | MAGIC | 42,854 | 17 | 0.010 | NA | NA | 87 | mmol/L | Scott et al., 2013 |
| Fasting glucose | MAGIC | 58,074 | 38 | 0.020 | NA | NA | 99 | mmol/L | Manning et al., 2012 |
| Fasting insulin | MAGIC | 51,750 | 38 | 0.020 | NA | NA | 99 | log pmol/L |  |

Characteristics of the consortia used in our study: number of samples, number of SNP included in the IV2 for different outcomes, proportion of phenotype variance explained by the instruments (tested in UKB), statistical power for a fixed effect of 0.15 SD (continuous traits) or 20% (binary traits) per SD change in BW, beta (continuous traits), OR (T2D) or HR (cardiovascular outcomes) from observational analyses in UKB and statistical power calculated for this observed association.
Abbreviations: BW, birthweight; CAD, coronary artery disease; AF, atrial fibrillation; IS, ischemic stroke; SBP, systolic blood pressure; UKB, UK Biobank; DBP, diastolic blood pressure; BMI, body mass index; WHR, waist-to-hip ratio; HDL, high density lipoprotein; LDL, low density lipoproteins; TG, triglycerides; T2D, type 2 diabetes; EGG, Early Growth Genetics; CAD, CARDIoGRAMplusC4D; AFGen, Atrial Fibrillation Genetics; ISGC, International Stroke Genetics Consortium; ICBP, International Consortium for Blood Pressure; GIANT, Genetic Investigation of ANthropometric Traits; GLGC, Global Lipids Genetic Consortium; DIAGRAM, DIAbetes Genetics Replication and Meta-analysis; MAGIC, Meta-Analysis of Glucose and Insulin related traits Consortium; SD, standard deviation.

### Statistical methods

#### Observational analysis

After confirming normal distribution of all continuous variables, we performed multivariable linear regression models to assess associations of BW with SBP, DBP, BMI, body fat, and WHR; and multivariable logistic regression models to study associations of BW with T2D and lipid medications. Multivariable-adjusted Cox proportional hazards models were performed to assess associations of BW with CAD, AF, IS, hemorrhagic stroke and heart failure events, separately; during a median follow-up time of 6.1 years (maximum 6.7 years). We use the DAGitty web tool (http://dagitty.net/dags.html) to systematically construct our multivariable model adjusting for confounders (Supplemental Figure I). All association analyses were adjusted for age, sex, region of the UKB assessment center, ethnicity, maternal smoking and Townsend index. We assessed evidence of nonlinear effects of BW on different outcomes using spline regression models. All observational analyses were performed in the UKB.

#### Mendelian randomization

We performed two-sample Mendelian randomization analyses using publically available consortia data, except for blood pressure where we performed a GWAS in UKB. We assessed the causal relationships of BW with CAD, AF, IS and T2D, and risk factors for these diseases (SBP, DBP, BMI, WHR, HDL, LDL, TG, 2- hour glucose, fasting glucose and fasting insulin) using the two-sample MR approach^19,20^. In order to minimize the risk of pleiotropy affecting our results, we performed analyses using three different IVs:

IV1) Including up to 58 independent lead variants (excluding the IGF2 locus due to imprinting; see Online Data Supplement) from the GWAS of BW performed by the EGG consortium^8^;
IV2) Including up to 46 variants after exclusion of 12 variants associated with CAD, AF, IS and T2D at GWAS significance; any confounders at GWAS significance; or with any of the confounders or CAD, AF, IS and T2D at a P-value lower than the P-value for association with BW (Supplemental Figure II). These associations were estimated in UKB.
IV3) Excluded 1-9 heterogenetic variants (different for each outcome; Supplemental Figure III). We performed a stepwise downward “model selection” in which SNPs were iteratively removed from the risk score until the heterogeneity test was no longer significant at the pre-specified threshold (*P*<0.05) using the R package *gtx.*

We decided *a priori* that IV2 would constitute our main model (balancing high statistical power and low risk of pleiotropy), but included IV1 to maximize power and IV3 to decrease risk of pleiotropy in sensitivity analyses.

We used four separate methods to estimate causal effects: the standard inverse-variance weighted (IVW) regression, the robust penalized IVW; as well as two robust regression methods, the weighted median-based method, and Egger regression^20^. We performed leave-one-out sensitivity analyses to identify if a single SNP was driving an association. To further address whether BW had a causal effect on CAD and T2D independently of BMI, we used a multivariate MR weighted regression-based method, in which the causal effects of multiple related risk factors can be estimated simultaneously^21,22^.

We estimated statistical power for the different MR analyses (Table 1) using sample sizes and variance explained specific for each analysis and an alpha threshold of 0.05 for two different effect sizes: 1) Assuming a fixed effect across phenotypes of 0.15 SD (continuous outcomes) or 20% (odds ratio, 1.2; dichotomous outcomes); and 2) For traits that were available in UKB, the effect size from observational analyses.
MR analyses were conducted with the R packages *TwoSampleMR* ^23^, and *MendelianRandomization* ^24^. Power for MR analyses was estimated with an online tool by Burgess (https://sb452.shinyapps.io/power/). Observational analyses were conducted with the R package *Survival* (version 3.3.0).

A flow chart of the different data sources used in this study is shown in Supplemental Figure IV. A detailed description of material and methods can be found in the Online Data Supplement.

## Results

In UKB, the mean age at baseline was 55.0 years (SD, 8.1 years) and 61% of subjects were females. During follow-up, 5,542 incident CVD cases occurred in participants free from the disease at baseline (2,656 CAD; and 1,580 AF; 688 IS; 363 hemorrhagic stroke; and 255 heart failure events; Supplemental Table I and II).

### Observational analyses

The results from observational analyses are summarized in Figure 1 (full results in Supplemental Table II). We observed strong inverse associations between BW and blood pressure, CAD, T2D and lipid-lowering treatment. In contrast, we observed strong and positive associations between BW and BMI and body fat percentage. After adjusting for multiple testing (12 traits), the associations were non-significant for WHR, AF, IS, hemorrhagic stroke and heart failure. We excluded non-linear associations between BW and any outcomes tested (*P*>0.05) by spline-regression (Supplemental Figure V).

**Figure 1.**
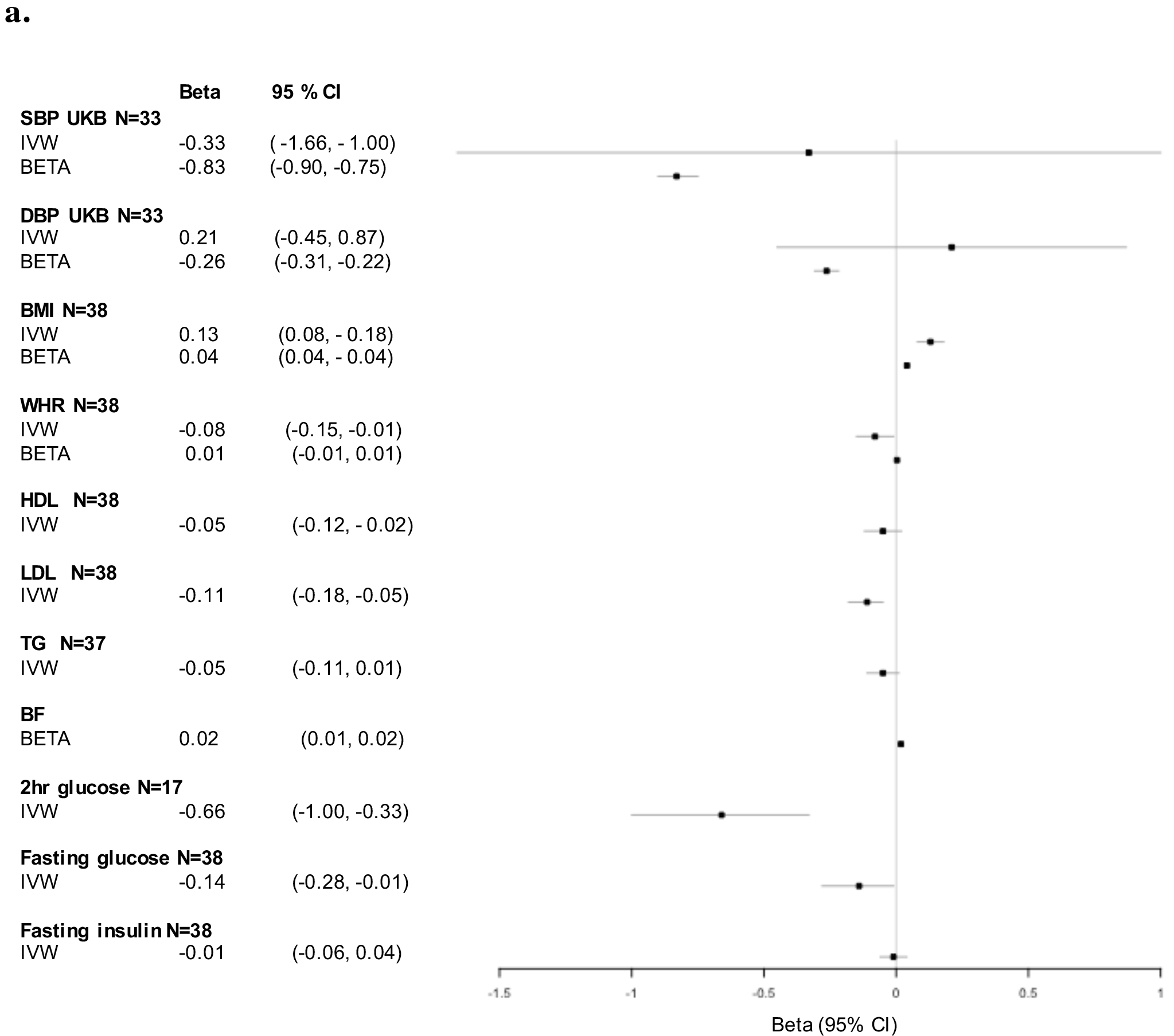
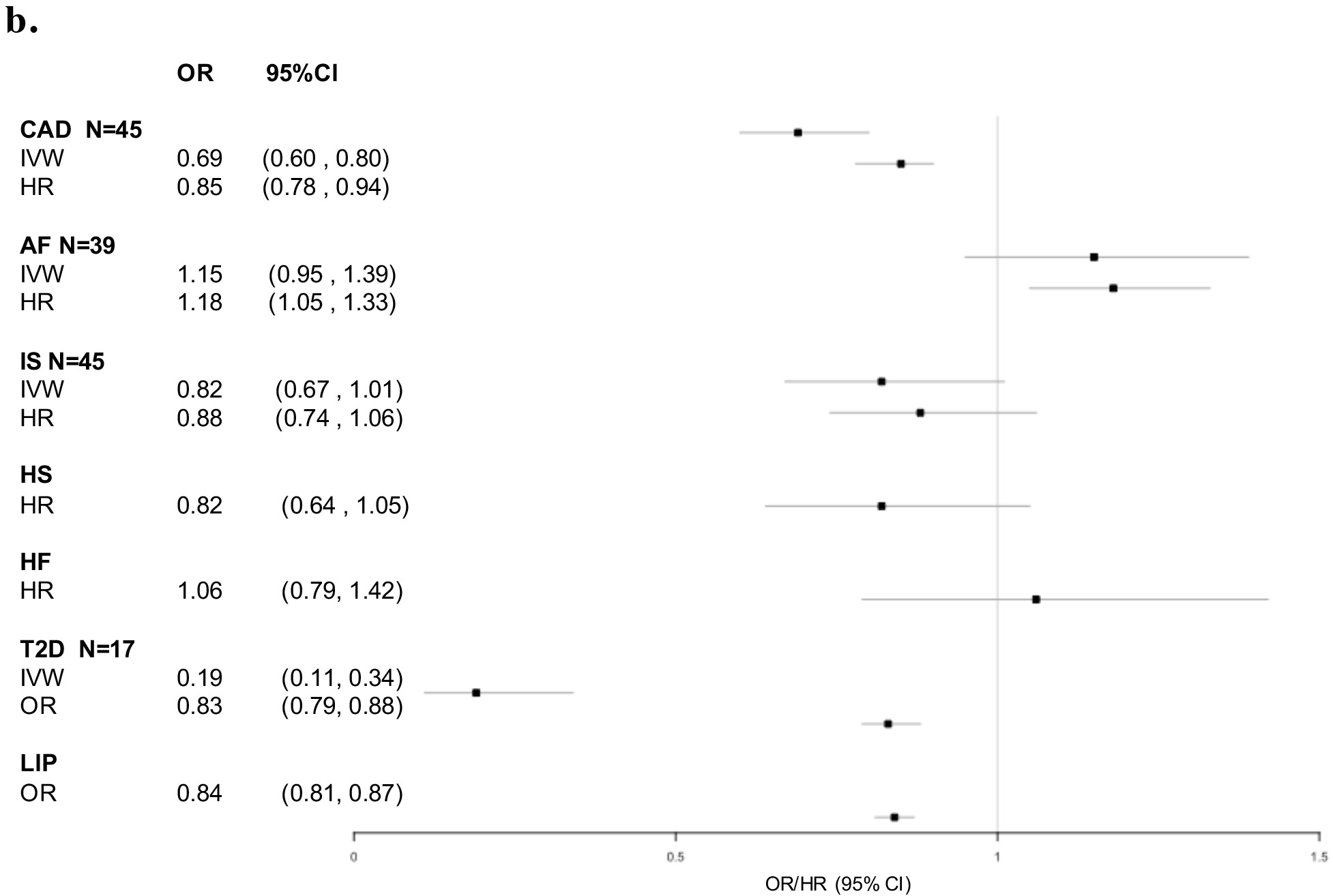
Inverse-variance weighted Mendelian Randomization (IVW) and association results (BETA/HR/OR) of birth weight (BW) with cardiovascular outcomes in UK Biobank using multivariable-adjusted linear and logistic regression, and multivariable-adjusted Cox proportional hazards models. **a.** Continuous outcomes: systolic and diastolic blood pressure in UK Biobank (SBP UKB, and DBP UKB; respectively), body mass index (BMI), waist-to-hip ratio (WHR), high density lipoprotein (HDL); low density lipoproteins (LDL); triglycerides (TG); body fat percentage (BF); 2-hour glucose; fasting glucose; and fasting insulin. **b.** Binary outcomes: coronary artery disease (CAD); atrial fibrillation (AF); ischemic hemorrhagic stroke (IS and HS, respectively); heart failure (HF); type 2 diabetes (T2D) and lipid medications (LIP).

[The betas from linear regression represent SD change in outcome variable per SD change in BW, except for SBP and DBP where they represent the outcome in raw unit (mmHg) per SD change in BW. Mendelian randomization analyses were based on the 46 variants included in the instrument variable 2 using data sources listed in Table 1. All effects for the IVW (beta or OR) are given in original units as provided by the consortia. Model adjustment: age, sex, region of the UKB assessment center, ethnicity, maternal smoking and Townsend index. Abbreviations: HR, hazard ratio; OR, odds ration; CI, confidence interval; N= number of variants included in the instrument variable.]

### Mendelian randomization

In our main analyses (IVW using the 46-SNP instrumental variable [IV2]), we found evidence of causal associations of BW with BMI, LDL, 2-hour glucose, CAD and T2D (Figure 1). The direction of the effect was negative for all the above outcomes (i.e. higher BW was associated with lower risk and vice versa), with the exception of BMI, where higher BW was associated with higher BMI. We did not find evidence of causal effect of BW on HDL, TG, fasting insulin, AF, and IS.

The leave-one out sensitivity analysis did not highlight any heterogeneous SNPs with a large effect on the results. After excluding heterogeneous SNPs in the IV3, our analysis showed no significant heterogeneity and no significant directional horizontal pleiotropy (all P>0.05; Supplemental Figure VI).

The analyses using penalized robust IVW, MR Egger, and weighted median methods consistently yielded similar effect estimates, but as expected with wider confidence intervals, especially for Egger regression (Supplemental Table III and Supplemental Figure VII). Further, sensitivity analyses using alternative IVs with higher power (IV1) and lower risk of pleiotropy (IV3) also provided similar results (Supplemental Table III).

The mediation analysis using the multivariate MR weighted regression-based method showed an independent association between BW and CAD, as well as between BW and T2D, not mediated by BMI in either case. The direction of the effect detected was consistent with our main MR analyses (Supplemental Table IV).

We had good statistical power to detect causal associations for all traits when assuming a fixed effect across phenotypes of 0.15 SD (continuous outcomes) or 20% (odds ratio, 1.2; dichotomous outcomes). When using the effect sizes from observational analyses of traits that were available in UKB, the power was adequate for all traits except DBP and WHR.

## Discussion

### Principal Findings

In this study of 237,631 individuals from the general population, we used self-reported BW as a proxy for fetal development to analyzed downstream consequences of intrauterine growth restriction. We describe the association of BW with incidence of T2D and five cardiovascular outcomes (CAD, AF, IS, hemorrhagic stroke and heart failure), and cardiometabolic risk factors (blood pressure, BMI, body fat, and WHR), and we identify a causal role of BW in the development of several cardiometabolic diseases. Our principal findings are several. First, in our observational study, we established that self-reported BW displays strong inverse associations with blood pressure, CAD and T2D, and strong direct associations with BMI and body fat. Second, our MR analyses indicate that low BW is causally related to higher risk of LDL and 2-hour glucose; and higher CAD and T2D in adults. This highlights the influence of prenatal determinants of fetal growth on the development of cardiometabolic diseases in adulthood. Third, our study suggests high BW to be causally associated with increased BMI, but not causally associated with blood pressure. Taken together and considering the different direction of the causality for BMI and CAD/T2D (higher BW increases BMI; lower BW increases CAD and T2D), our results suggest a causal association of intrauterine growth restriction and low BW with risk for CAD and T2D, an association which does not appear to be mediated by obesity or hypertension.

In their initial description of the *“thrifty phenotype hypothesis”*^3^, Barker and Hales proposed that BMI would be a possible mediator of the associations detected between low BW and adult T2D and CAD. The hypothesized primary effect of BMI was supported by evidence from both population and experimental studies linking low BW with predisposition to an increased risk of metabolic diseases such as T2D^25,26,27,28,29^, hypertension^30,31^ and CAD^32^. However, in our study and in prior observational analyses, *higher* BW is associated with obesity (a universally recognized correlate of cardiometabolic disease) in both childhood^33,34^ and adulthood^35,8^. Our findings suggest a causal association of low BW with CAD and T2D, which is uniquely independent of the relationship between high BW and increased BMI. Consistent with our observed effects of low BW on risk for CAD and T2D independent of adult obesity, a recent study of African American women failed to detect a causal role for BMI in mediating the increased risk for T2D in adult life among individuals with low BW^36^. New models for how risk for cardiometabolic disease in adulthood is directly conferred by growth restriction *in utero* without a compensatory change in BMI are needed to explain our observation of a direct causal relationship.

Explicit in the Barker hypothesis and explored by the experimental literature^37,38^, is a model in which prenatal growth stress leads to metabolic reprogramming beginning *in utero*. In the setting of prenatal malnutrition, the fetus is hypothesized to shift toward insulin resistance in order to allow for maximum uptake of available energy and nutrients. In this hypothesis, the persistence of insulin resistance after parturition might then trigger rapid postnatal growth with the concomitant potential for increased long-term risk of T2D, obesity and CAD in adulthood^25,39^. However, our findings support a separate direct causal link between intrauterine growth restriction and long-term risk for cardiometabolic disease, which does not involve adult obesity. Consistent with our detection of a causal relationship, one prior report using IV analyses, but with much fewer variants also described a direct causal association between low BW and T2D^5^.

In contrast to our results, Yeung et al.^6^, reported no causal association between BW and CAD. However, this study was based on a weak instrumental variable consisting of seven SNPs, explaining only 0.45% of the variance for BW (in contrast with our score that explained 2.2% of the variance), resulting in limited statistical power of 56% suggested by *post-hoc* calculations. In this context, it is also worth mentioning the genetic correlation analyses of BW with several health-related traits, published in the recent GWAS for BW used to create IVs for our MR study^8^. As in our study, they reported strong positive genetic correlations with BMI, and inverse genetic correlations with CAD and T2D. In contrast to our MR results, they highlight a negative genetic correlation with SBP. This discrepancy is probably related to the different methods used. Indeed, they used the linkage-disequilibrium score regression model^40^ which use all GWAS summary statistics of the traits of interest to estimate the genetic correlations, while MR methods are based on a much smaller number of variants, aiming to decrease the risk of horizontal pleiotropy driving associations.

### Clinical Implications

Given our observation that low BW is causally related to LDL, 2-hour glucose, CAD and T2D, these findings are strongly consistent with the growing recognition of the long-term public health importance of supporting adequate prenatal nutrition. Diet is a broadly modifiable risk factor, and both maternal and paternal nutrition have an impact on the risk of metabolic syndrome, lipid dysregulation, fat deposition, obesity and hypertension in offspring via a hypothesized mechanism of *in utero* epigenetic imprinting^41,42,43^. Both epidemiological and animal studies highlight that undernutrition, overnutrition, and inadequate diet composition negatively impact fetoplacental growth and metabolic patterns, potentially having adverse later life metabolic effects for the offspring^44^. Additionally, our data may also offer a window into the role by which non-nutritional factors affecting fetal growth such as congenital heart disease and premature birth, may predispose affected individuals to long-term risk of cardiometabolic disease in adulthood^45,46,47^.

Our results indicate that some proportion of common chronic diseases of adulthood could potentially be reduced by achieving optimal fetal nutrition. Short-term follow-up of children born after randomized nutritional interventions in pregnancy describe beneficial effects on growth, vascular function, lipid levels, glucose tolerance and insulin sensitivity; though longer-term studies examining nutrition and growth in premature infants display a more complex set of relationships^48,49^. Considered in the context of populations, our data suggest that attention to prenatal nutrition and intrauterine growth may have long-term consequences regarding the risk of CAD, obesity and diabetes in adult life.

### Strengths and limitations

To our knowledge, this is the largest and most comprehensive study of associations of BW with outcome to date. Additionally, we used three different IVs to maximize power and to decrease risk of pleiotropy, and several methods for MR analyses all yielding consistent effects for the tested hypotheses. However, our study is limited by the study samples of middle-aged to elderly individuals of European descent from rich countries. Hence, generalizability of our findings to other populations where the diet, prenatal care, prevalence and predispositions of cardiometabolic disease are different is unknown. Further, although we excluded variants with higher likelihood of pleiotropy from our analysis and applied a range of sensitivity analyses and methods robust to pleiotropy, little is known about the mechanisms underlying loci included in the IV. Though our comprehensive analytical framework did not indicate any presence of horizontal pleiotropy, it is possible that some or all of these loci may also have a direct influence on the processes leading to CAD or T2D independent of intrauterine growth. Finally, despite the large sample in this study, statistical power to detect potentially causal relationships was limited for some traits, at least for the effect sizes from our observational analyses (in particular, DBP and WHR; Table 1).

### Conclusion

In conclusion, we demonstrate that intrauterine growth restriction, as evidenced by lower BW, is causally related with increased susceptibility to T2D and CAD, but that this effect is independent of adult hypertension or obesity, which has been previously hypothesized to be mediator of such association. Our study supports the notion that population level interventions improving prenatal nutrition and growth may improve cardiometabolic disease profiles later in life, but this needs to be confirmed using other study designs, such as large-scale community-based intervention trials.

## Acknowledgements

This research has been conducted using the UK Biobank Resource under Application Number 13721. Data on BW; CAD; AF; IS; SBP and DBP; BMI and WHR; HDL, LDL, and TG; T2D; 2-hour glucose, fasting glucose and fasting insulin have been contributed by EGG; CARDIoGRAMplusC4D; AFGen; ISGC; ICBP; GIANT; GLGC; DIAGRAM and MAGIC investigators, respectively.

